# Resveratrol reduces COMPopathy in mice through activation of autophagy

**DOI:** 10.1101/2020.10.26.355628

**Authors:** Jacqueline T. Hecht, Francoise Coustry, Alka C. Veerisetty, Mohammad G. Hossain, Karen L. Posey

## Abstract

Misfolding mutations in cartilage oligomeric matrix protein (COMP) cause it to be retained within in ER of chondrocytes, stimulating a multitude of damaging cellular responses including ER stress, inflammation and oxidative stress which ultimately culminates in the death of growth plate chondrocytes and pseudoachondroplasia (PSACH). Previously, we demonstrated that an antioxidant, resveratrol, substantially reduces the intracellular accumulation of mutant COMP, dampens cellular stress and lowers the level of growth plate chondrocyte death. In addition, we showed that resveratrol reduces mTORC1 (mammalian target of rapamycin complex 1) signaling, suggesting a potential mechanism. In this work, we investigate the role of autophagy in treatment of COMPopathies. In cultured chondrocytes expressing wild type or mutant COMP (MT-COMP), resveratrol significantly increased the number of large LC3 vesicles, directly demonstrating that resveratrol stimulated autophagy is an important component of the resveratrol-driven mechanism responsible for the degradation of mutant COMP. Moreover, pharmacological inhibitors of autophagy suppressed degradation of MT-COMP in our established mouse model of PSACH. In contrast, blockage of the proteasome did not substantially alter resveratrol clearance of mutant COMP from growth plate chondrocytes. Mechanistically, resveratrol increased SIRT1 and PP2A expression and reduced MID1 expression and activation of pAKT and mTORC1 signaling in growth plate chondrocytes, allowing clearance of mutant COMP by autophagy. Importantly, we show that optimal reduction in growth plate pathology, including decreased mutant COMP retention, decreased mTORC1 signaling and restoration of chondrocyte proliferation was attained when treatment was initiated between birth to one week of age in MT-COMP mice, translating to birth to approximately 2 years of age in PSACH children. These results clearly demonstrate that resveratrol stimulates clearance of mutant COMP by an autophagy-centric mechanism.

## Introduction

Next generation sequencing has resulted in a huge expansion of information that has led to greater understanding of the mutational underpinnings of genetic disorders^(1)^. This is particularly true for skeletal dysplasias, in which 425 distinct disorders are caused by mutations in 437 different genes^(1)^. Mouse modeling of the genetic mutations generating specific skeletal dysplasia has revealed the mechanistic pathways that underlie the resulting pathologies, thereby identifying therapeutic targets. Previously, we showed that mutations in cartilage oligomeric matrix protein (COMP) causes pseudoachondroplasia (PSACH; COMPopathy), a well-recognized and clinically characterized severe dwarfing condition^(2)^. The MT-COMP transgenic mouse, with the common D469del mutation that recapitulates the short stature and skeletal finding of PSACH, was used to define the pathology that specifically affects growth plate chondrocytes^(3–7)^. In this study, mechanistic-driven therapeutic approaches in the MT-COMP mouse were used to define the mechanisms of resveratrol therapy.

Chondrocytes are secretory cells that generate abundant extracellular matrix proteins necessary to build cartilage. COMP is normally synthesized in the rER cisternae and is processed through the Golgi then exported to the ECM where it forms an ordered matrix network^(8)^. It has long been known that mutant protein accumulates in PSACH chondrocytes^(9–11)^, mutations in COMP were shown to cause PSACH^(10,11)^. It was subsequently shown that mutant COMP misfolds in the ER and prematurely assembles an ordered matrix involving types 2 and 9 collagens, matrilin 3 and other extracellular proteins, resulting in massive intracellular protein accumulation^(7,12,13)^. The protein accumulation in the ER of chondrocytes initiates a self-perpetuating pathological loop between oxidative stress, inflammation and ER stress that leads to elevated mTORC1 signaling, DNA damage and death of chondrocytes^(4–6,14)^. Misfolded proteins in the ER are typically degraded through macroautophagy (henceforth referred to as autophagy) or by the proteasome after being translocated out of the ER^(15)^. In the presence of mutant COMP, the high volume of stalled protein/mis-folded protein/intracellular matrix either overwhelms the cellular clearance mechanisms and/or blocks export and degradation. In either situation, these clearance mechanisms do not efficiently clear mutant COMP from the ER and normal chondrocyte function is disrupted.

Previously, we demonstrated that treatment of MT-COMP mice with resveratrol or aspirin from birth to 4 weeks decreased mutant COMP intracellular retention, inflammation, chondrocyte cell death and restored chondrocyte proliferation^(3,4,7,16)^. Notably, these treatments rescued 50% of the loss of femur length. These studies now define the cellular processes stimulated by resveratrol (aspirin) that prevent mutant COMP accumulation and, importantly, define the appropriate timing of treatment.

## Material and Methods

### Bigenic mice

The MT-COMP (C57BL/6 mus musculus) mice used in these and previously described experiments contain the pTRE-COMP (coding sequence of human COMP+FLAG tag driven by the tetracycline responsive element promoter) and pTET-On-Col II (rtTA coding sequence driven by a type II collagen promoter)^(17–19)^. The presence of these two transgenes had not led to any readily detectable health concerns. Male and female mice were administered DOX (500 ng/ml) postnatally in their drinking water until collection. Mice were housed as a single sex group after weaning and fed standard chow (PicoLab rodent diet 20 #5053). Ten mice were in each treatment group or control group and litters were assigned to each group until each group contained at least 10 animals. Investigators were not blinded during allocation and animal handling but were blinded at analysis. These studies were approved by the UTHealth Animal Welfare Committee.

### Immunohistochemistry

Hind limbs from male MT-COMP and C57BL\6 control mice were collected and tibial growth plates analyzed as previously described ^(52)^. The limbs were fixed in 95% ethanol and pepsin (1mg/ml in 0.1N HCl) was used for antigen retrieval. Immunostaining was performed by incubating different sections overnight at 4 °C using the following antibodies: human COMP (Abcam Cambridge, MA ab11056-rat 1:100), pS6 (1:200 2215S rabbit polyclonal Cell Signaling Technology), PI3K-1 (abcam-ab 225720-1:200), SIRT-1 (abcam: ab 32441 - 1:100), pAMPK (Invitrogen: PA537821-1:100), interleukin 16 (IL-16) (Santa Cruz Biotechnology; sc-7902, 1:100), YM1/ECF-L (Stem Cell Technologies; 01404, 1:100) and PCNA (abcam ab92552 - 1:200). Species specific biotinylated secondary antibodies were used for 1 hr at RT. Sections were then washed and incubated with streptavidin horseradish peroxidase (HRP) and DAB was used as chromogen. The sections were dehydrated and mounted with Thermofisher cytoseal 60 and then visualized under the BX51 inverted microscope (Olympus America, Center Valley, PA). Limbs were fixed in 10% wt/vol formalin for terminal deoxynucleotidyl transferase–mediated deoxyuridine triphosphate-biotin nick end labeling (TUNEL) staining. *A priori* data indicate that the percentage TUNEL positive chondrocytes are approximately 80% in untreated mice and 20% in mice treated with resveratrol from birth to 4 weeks, with within group standard deviations of around 12%. For a one-way ANOVA analysis with post-hoc Tukey test to identify difference in TUNEL percentages between 5 groups at a Type I error rate of 0.05, a sample size of 10 mice per group would have 80% power to identify an effect size (Cohen’s d) of 0.51. This would translate into pair-wise differences in means of at least 17% in at least two of the 5 groups.

### Drug treatments

No adverse events were associated with these drugs administered as described. Rapamycin treated 2 mg/kg reduced viability of pups and the dosage was reduced to 1 mg/kg.

#### Resveratrol

Resveratrol (Reserveages Organics liquid resveratrol super berry extract), with known antioxidant/anti-inflammatory properties^(20–22)^, was administered in a dose of 0.25 g/L in the drinking water^(3)^ starting either from birth (transmitted through maternal breast milk) or 1 or 2 or 3 weeks and continued until 4 weeks of age^(3)^.

#### Aspirin

Aspirin, a COX2 inhibitor that reduces inflammation was administered in the drinking water in a dose of 0.3 g/L of aspirin (Sigma, USA) from birth to 4 weeks^(3)^.

#### Rapamycin

Rapamycin (Tocris Bioscience, United Kingdom) inhibits mTOR pathway, inducing autophagy^(23)^, was dissolved in ethanol at 1 mg/ml and stored in −20°C. On the day of injection, rapamycin was diluted with PBS to 0.1 mg/ml. Intraperitoneal injection of rapamycin at 1 mg/kg was performed 5 days per week from 1 - 4 weeks of age^(24)^.

#### Metformin

Metformin (Sigma, USA) activates AMPK, which in turn represses mTORC1 activity stimulating autophagy^(25,26)^, was administered at 0.5 mg/ml in DOX drinking water from birth (through maternal breast milk) to 4 weeks ^(24,27)^.

#### Bafilomycin A

Bafilomycin A (Cayman Chemical, USA) blocks autophagy by preventing fusion of autophagosome with lysosome^(28)^ was dissolved in 10 mg/ml DMSO, aliquoted and stored at - 20° C. For each subcutaneous injection, bafilomycin A was freshly diluted to 0.05 mg/ml in PBS. Mice were injected with 0.3 mg/Kg of bortezomib 5 times per week beginning at age 1 week^(29)^. Bafilomycin A administration to untreated MT-COMP did not alter mutant COMP accumulation.

#### Bortezomib

Bortezomib (# 2164 AvaChem Scientific, USA) inhibits the chymotryptic activity of the proteasome (20S subunit) reducing proteasomal degradation^(30)^ was dissolved in 10mg/1ml DMSO, aliquoted and stored at −20° C. Freshly diluted bortezomib was diluted for each subcutaneous injection to 0.01 mg/ml in PBS. Mice were injected with 0.1mg/Kg of bortezomib 3 times per week beginning at age 1 week^(31,32)^. Bortezomib administration to untreated MT-COMP did not alter mutant COMP accumulation.

### Cell culture

RCS cells expressing either human wild-type or MT-COMP were generated using the Lenti-X-Tet-On advanced inducible expression system according to the manufacturer protocol (Takara Bio Company, Mountain View, CA). Briefly, RCS cells were infected with high-titer lentiviral preparations of pLVX-Tet-on advanced vector or pLVX-Tight-human D469del-MT-COMP vector; and selected with puromycin and G418. Expression of human COMP in RCS cells was validated by Western Blot with anti-Flag antibody to recognize the tagged COMP (1:5000 F7425 rabbit polyclonal from Sigma, Saint Louis, MO). DOX was used for 4 days to induce COMP expression in RCS cells, which were then treated for 1 day as follows: 1) control with no chloroquine and no resveratrol, 2) chloroquine (50 μM), 3) resveratrol (50 μM) and 4) both resveratrol (50 μM) and chloroquine (50 μM). The cells were fixed post-treatment with methanol at −20°C for 10 minutes, rinsed with PBS and stored at 4°C. Coverslips were then incubated for 1 h at 37°C with LC3 primary antibody (Cell signaling; 2775; 1:500). Primary antibodies were detected using Alexa Fluor 594 for 1 h at 37°C (dilution 1:1000). Images were collected with confocal microscope at 63x oil immersion lens (Zeiss LSM-510). *A priori* data suggests that the median number of foci per image for each of the 8 groups would range from 0 through 2 with interquartile ranges of 0 through 3. Assuming a power of 0.80 and a Type I error rate of 0.05, a sample size of 23 cells counted per image will provide us with sufficient power to identify a small effect size of 0.1 (Cohen’s d). For our analysis, we elected to count 30 to 50 cells per image. ImageJ was used to count LC3 foci on ten images for each group with 30-50 cells per image. T-tests were used to compare each group to the other groups and scorers were blinded.

## Results

### Resveratrol treatment reactivates autophagy in MT-COMP chondrocytes

Autophagy and proteasomal degradation were examined as they are the cellular mechanisms responsible for removing misfolded protein from the ER^(33)^. Previously, we showed that MT-COMP accumulation in the ER of murine MT-COMP growth plate chondrocytes occurs because autophagy was blocked by elevated mTORC1 signaling^(4)^. It remains unclear whether proteasome degradation plays a role in this ER-stress-induced chondrocyte pathology because COMP-ECM matrix within the ER may prohibit translocation from the ER to the proteasome ^(4)^. To determine whether autophagy and/or the proteasomal degradation were responsible for resveratrol clearance of MT-COMP, neonatal mice were treated with resveratrol birth to 4 weeks in the absence or presence of an autophagy blocker (bafilomycin A^(28)^) or a proteasomal inhibitor (bortezomib^(30)^) from 1 to 4 weeks of age. As shown in **Figure 1 Panel 1B**, resveratrol treatment alone dramatically reduced MT-COMP accumulation compared to untreated growth plate chondrocytes (**Fig. 1 Panel 1A**)^(3)^. The addition of bafilomycin A largely prevented resveratrol-stimulated clearance of MT-COMP (**Fig. 1 Panel 1C**) while bortezomib only partially reduced MT-COMP clearance (**Fig. 1 Panel 1D**). This shows that autophagy is the primary mechanism of resveratrol stimulated MT-COMP clearance and proteasome activity plays a minor role.

**Figure 1.**
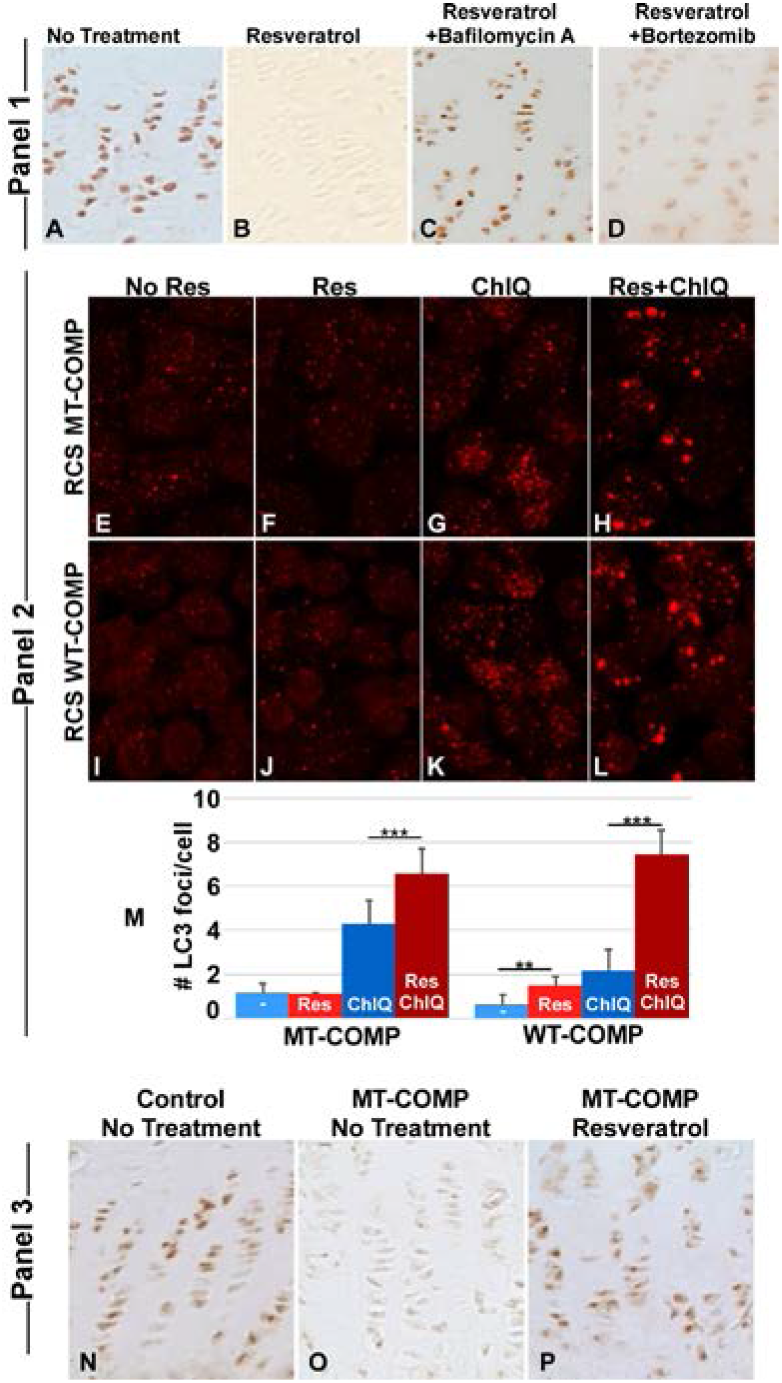
Reactivation of autophagy correlates with reduction in intracellular COMP retention. **Panel 1:** Growth plate sections from MT-COMP mice at age 4 weeks were immunostained with human COMP antibody after different treatments: **A)** no treatment, **B)** resveratrol treatment of 0.25 g/L in drinking water from birth to 4 weeks, **C)** resveratrol and bafilomycin A (0.3 mg/Kg injected 5X per week from 1-4 weeks) treatments or **D)** bortezomib (0.1mg/Kg 3X per week from 1-4 weeks). Bafilomycin A blocks the fusion of the autophagosome to the lysosome^(28)^, while Bortezomib is a proteasomal inhibitor ^(30)^. Bafilomycin A blocks autophagy in resveratrol treated growth plates (C) while bortezomib only partially blocks resveratrol (D) clearance of MT-COMP. Each treatment group included 10 animals. **Panel 2:** RCS cells expressing either WT- or MT-COMP immunostained for LC3: untreated (**E, I**), resveratrol (**F, J**), chloroquine (**G, K**) and resveratrol and chloroquine (**H, L**) treatments. Quantification of LC3 positive foci (**M**) in MT- and WT-COMP RCS cells. Bars = means with standard deviation. Chloroquine blocks fusion between the lysosome and autophagasome ^(28)^. Res = resveratrol; ChlQ = chloroquine; MT-COMP = mutant COMP; WT-COMP = wild-type COMP; **P ≤0.005; *** P ≤0.0005. LC3 foci were quantified in 30-50 cells in 10 different images for each group. **Panel 3:** SIRT1 immunostaining of control (no treatment) (**N**), untreated MT-COMP (**O**) and resveratrol treated MT-COMP (**P**) growth plates are shown. Ten animals were included in each treatment group

To further demonstrate that resveratrol stimulates autophagy in chondrocytes, resveratrol-induced autophagic vesicle clearance was pharmacologically blocked using chloroquine to allow visualization of autophagic vesicles, since they are rapidly degraded under normal conditions. Rat chondrosarcoma (RCS) cells expressing either wild type (WT) or mutant COMP were treated with resveratrol in presence or absence of chloroquine and the number of microtubule-associated protein light chain 3 (LC3) positive vesicles (foci), the classic marker of autophagy, were assessed^(4)^. As shown in **Fig. 1 Panel 2 E, F, I, J**, autophagy was not observed in the absence of chloroquine, as expected. The number of autophagic vesicles was significantly increased with resveratrol treatment (**Fig. 1 Panel 2 H, L, M**) over the basal level of autophagy (**G, K**) confirming that resveratrol stimulates autophagy in chondrocytes.

### Resveratrol promotes MT-COMP growth plate chondrocytes survival through SIRT1 signaling

Studies have shown that resveratrol increases SIRT1, promoting cellular survival and decreasing inflammation and oxidative stress^(22,35–40)^. SIRT1 enhances cellular survival by deacetylating LC3, which promotes autophagy^(22)^ To investigate the role of SIRT1 in resveratrol autophagy reactivation, growth plates were evaluated for the presence of SIRT1 in untreated MT-COMP mice and those treated from birth to 4 weeks of age with resveratrol. SIRT1 was suppressed in MT-COMP growth plate chondrocytes (**Fig. 1 Panel 3O**) and resveratrol treatment reverses the mutant COMP suppression of SIRT1 (**Fig. 1 Panel 3P, N**). These results demonstrate that resveratrol treatment promotes autophagy through SIRT1 in MT-COMP mice.

### Resveratrol modulates mTORC1 pathway signaling components

We have previously shown that AKT is activated by MID1 and suppressed by PP2A in the mutant COMP pathology^(4)^. AKT, an upstream regulator of mTORC1, is up-regulated by PI3K-I^(41,42)^. These studies evaluated whether PI3K-I elevates mTORC1 in MT-COMP growth plate chondrocytes in mice treated from birth to 4 weeks^(4)^. Shown in **Figure 2 Panel 1**, PI3K-I was absent in untreated MT-COMP mice compared to controls. The absence of PI3K-1 in untreated MT-COMP growth plate chondrocytes in which mTORC1 signaling is high^(4)^ indicates that PI3K-I does not play a role in elevating mTORC1 signaling and blocking autophagy in MT-COMP growth plate pathology.

**Figure 2.**
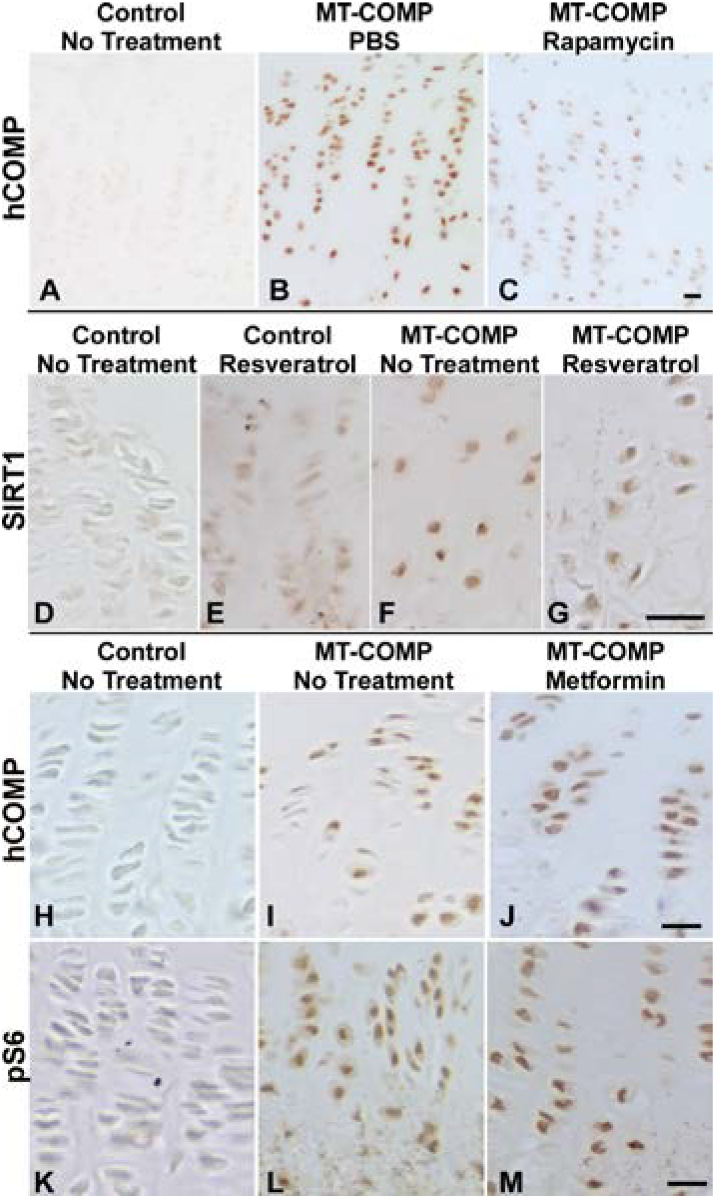
PI3K-1 and AMPK do not play a role in mutant COMP mTORC1 signaling. **Panel 1** shows results of resveratrol and **Panel 2** metformin treatments. Each treatment group contained 10 mice. Growth plates at 4 weeks: Control (**A, D, G, J**), MT-COMP untreated (**B, E, H, K**), resveratrol (0.25 g/L of resveratrol in drinking water from birth to 4 weeks) (**C**) or metformin (0.5 mg/ml in drinking water from birth to 4 weeks) (**F, I, L**). Immunostaining of PI3K-I (**A-C**), human COMP (**D-F**), pAMPK (**G-I**) and phospho-S6 (pS6) (**J-L**) is shown. Bars = 100 μm.

### Metformin does not activate autophagy clearance of MT-COMP

Metformin modulates mTORC1 signaling by activating adenosine monophosphate-activated protein kinase (AMPK) ^(43)^, which in turn dampens mTORC1 signaling stimulating autophagy ^(42,44)^. Previously, we showed that elevated mTORC1 signaling results from TNFα/TRAIL inflammation and ER stress^(4)^. We asked whether metformin treatment could induce autophagy in MT-COMP mice, thereby mimicking resveratrol’s mechanism of action. Here, we assessed whether mTORC1 signaling could be reduced by metformin-stimulated AMPK activation in growth plate chondrocytes. As shown in **Fig. 2 Panel 2,** metformin administered to MT-COMP mice from birth to 4 weeks increased AMPK levels as expected, indicating that metformin reached the growth plate in therapeutic concentrations. However, metformin did not decrease either mutant COMP retention or pS6, which is a measure of mTORC1 signaling. These results demonstrate that metformin did not repress mTORC1 signaling (as measured by pS6) or induce autophagy in MT-COMP mice indicating that metformin does not mimic resveratrol’s mechanism of action.

### Resveratrol postnatal treatment windows

Previously, we demonstrated that resveratrol treatment in MT-COMP mice from birth to 4 weeks of age dampens mutant COMP intracellular ER retention, inflammation, chondrocyte death, and restored chondrocyte proliferation, resulting in increased femoral length^(45)^. To define the treatment windows in which resveratrol could be administered to MT-COMP mice to reduce chondrocyte pathology, resveratrol was started at birth, 1, 2 or 3 weeks of age and growth plates were collected at 4 weeks. As shown in **Figure 3**, resveratrol treatment for 1 - 3 weeks diminished intracellular retention, inflammation and chondrocyte cell death. The most dramatic reduction of inflammatory markers, interleukin 16 (IL-16) and YM1 (YM1 also known as eosinophil chemotactic factor-lymphocyte (ECF-L)), was observed in growth plates treated for three or more weeks with resveratrol (**Fig. 3G-R**). Importantly, resveratrol treatment beginning as late as 3 weeks of age reduced inflammation and chondrocyte death (**Fig. 3I, O**). Decreased cell death (TUNEL positive chondrocytes) showed a significant temporal response with treatment (**Fig. 3A-F; M; EE)**. In contrast, chondrocyte proliferation did not recover until 3 weeks of resveratrol treatment (**Fig. 3Y-DD; FF)**. Importantly, the timing of autophagy derepression (**Fig. 3EE-JJ** pS6) correlated with recovery of chondrocyte proliferation in response to resveratrol treatment suggesting that appropriate autophagy function plays a role in resveratrol’s therapeutic action and growth plate chondrocyte cellular health.

**Figure 3.**
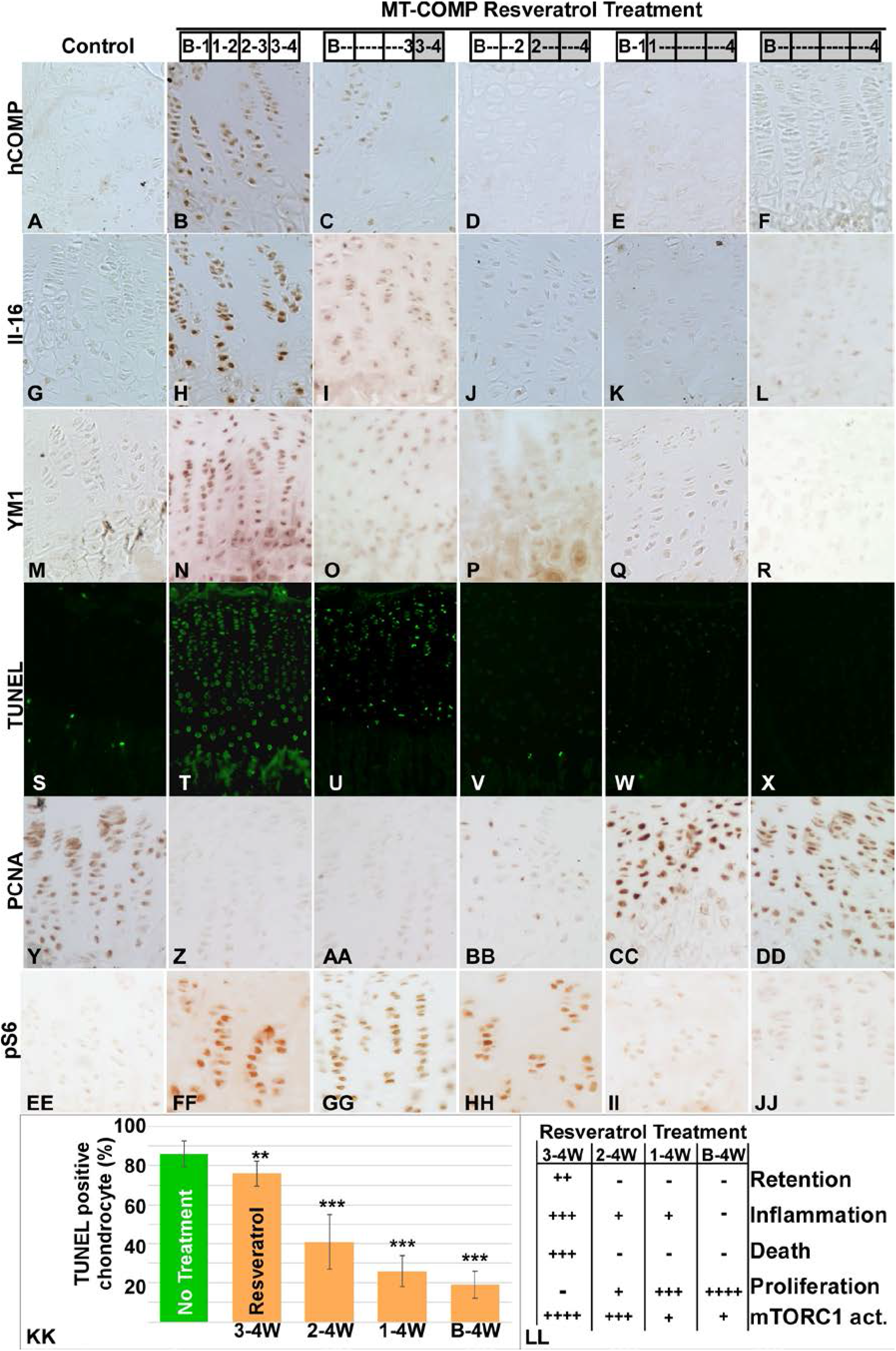
Resveratrol treatment window. Treatments started at 1 or 2 or 3 weeks after birth and stopped at 4 weeks and evaluated for mutant COMP pathology. Ten mice were included in each treatment group. T reatments starting at birth or 1 week of age produced the best therapeutic outcomes (**K, L**). Later therapy dampened but did not eliminate the disease progression and did not restore DNA proliferation (**C**). Resveratrol treatment times are shown in the shaded boxes. Control (C57BL\6) (**A, G, M, S, Y, EE**) and MT-COMP with no treatment (**B, H, N, T, Z, FF**) resveratrol treatment for 1 (**C, I, O, U, AA, GG**), 2 (**D, J, P, V, BB, HH**), 3 (**E, K, Q, W, CC, II**) or 4 (**F, L, R, X, DD, JJ**) weeks and growth plates at 4 weeks were evaluated for human COMP retention, IL-16 and YM1, TUNEL (cell death), PCNA (cell proliferation) and mTORC1 signaling (pS6). Quantification of TUNEL positive chondrocytes for different treatment periods are shown in **KK** and summary of findings in **LL**. Bars in KK = means with standard deviation. Resveratrol treatment decreased intracellular MT-COMP (**C**) and IL-16 and YM1 inflammation (**I, O**) after one week of treatment compared to untreated MT-COMP growth plates (**H, N**) and mTORC1 signaling was substantially reduced after 3 weeks of treatment (**II, JJ**). The relative change of MT-COMP pathology on treatment period is shown in **LL**.

## Discussion

The results of these studies establish that obstruction of autophagy causes mutant COMP accumulation in the ER of growth plate chondrocytes and, importantly, resveratrol clears and/or prevents intracellular retention by reactivating and/or maintaining autophagy function. The mechanism underlying this COMPopathy is the culmination of many experimentiments^(3,5,46)^, is caused by the retention of mutant COMP in the ER, which stimulates a chronic self-perpetuating pathological loop between ER stress, inflammation and oxidative that elevates mTORC1 signaling through a complex network involving CHOP and TNFα^(4)^ (**Fig. 4A**). This this pathology can be interrupted by the loss of CHOP^(5,46)^ demonstrating that CHOP is an important therapeutic target. Resveratrol’s multifocal targeted mechanisms of action is shown in **Figure 4B** includes the repression of CHOP.

**Figure 4.**
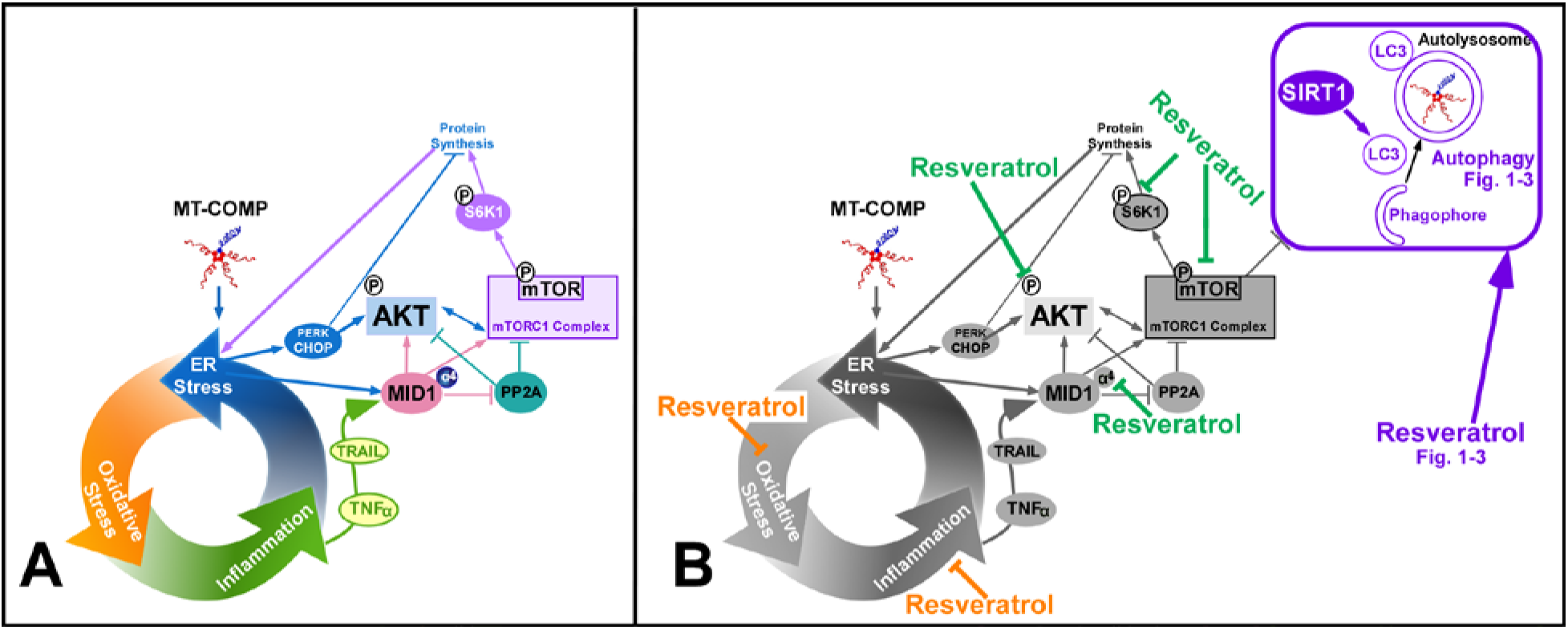
Mechanisms by which resveratrol treatment inhibits MT-COMP growth plate chondrocyte pathology. **A)** Pathologic mechanism initiated and maintained by retention of MT-COMP in ER of growth plate chondrocytes of MT-COMP mice^(3–5)^. **B)** Resveratrol actions in pathologic processes that reduced chondrocyte ER stress including less inflammation and oxidative stress^(3)^ (shown in orange) and decreases pAKT, MID1 and mTORC1 signaling which blocks autophagy^(4)^ (green) through activation of SIRT1 that promote autophagosome maturation (shown in purple).

Our previous studies showed that resveratrol and aspirin dramatically reduce intracellular mutant COMP in the ER of MT-COMP growth plate chondrocytes, but the mechanism(s) is unknown^(3)^. *In vitro* and *in vivo* approaches were used to identify and validate the mechanism by which resveratrol reduces the ER intracellular retention. RCS cells expressing either mutant or wildtype COMP showed that autophagy was increased when treated with resveratrol (**Fig. 2)** and rapamycin treatment, an autophagy-inducing drug ^(47)^, decreased the intracellular mutant COMP accumulation in growth plate chondrocytes in MT-COMP mice (**Supplementary Fig. 2**). Furthermore, pharmacologically blocking autophagy inhibited resveratrol clearance of mutant COMP demonstrating that resveratrol stimulated autophagy is the key molecular mechanism that clears mutant COMP preserving chondrocyte function and preventing cell death in MT-COMP mice (**Fig. 1**). These findings are consistent with previous studies in other cell types which showed that autophagy is the primary mechanism utilized to clear protein aggregates from the ER ^(48)^.

Autophagy plays an essential role in cellular maintenance by recycling cellular contents and participating in ER protein quality control^(49)^. Many storage disorders involve a failure of or inefficient autophagy including proteinopathies (Alzheimer disease, tauopathies, Parkinson disease, Huntington disease), lipid storage disorders (Niemann Pick type C disease, Gaucher disease, Fabry disease) and glycogen storage disorders (Lafora disease, Von Gierke disease, Pompe disease)^(50,51)^. Autophagy failure in disease occurs through a variety of mechanisms including impaired autophagosome formation, compromised autophagosome maturation (through mTOR activation, AMPK inhibition, SIRT1 downregulation) and improper recognition of autophagic cargo^(51)^. In PSACH, a COMPopathy/proteinopathy, mTORC1 activation and SIRT1 downregulation blocks autophagosome maturation through a complex network (**Figs. 1** and **4**)^(4)^. Chronic inflammation has been shown to reduce SIRT1 expression^(52–56)^ and the persistent inflammation in MT-COMP growth plate chondrocytes is likely to account for SIRT1 repression in MT-COMP mice^(3)^. Recently, we showed that ER stress and TNFα/TRAIL upregulated MID1 and MID1 increased mTORC1 signaling directly and by decreasing PP2A and increasing phosphoAKT^(4)^. TNFα/TRAIL upregulation of MID1 has only been observed in airway inflammation and fibrosis^(57–60)^. This link between ER stress and MID1 is novel to the mutant COMP pathology^(4)^ (**Fig. 4A**). In contrast, PI3K-I and AMPK do not play a role in elevation of mTORC1 signaling in mutant COMP growth plate chondrocyte pathology (**Fig. 2**), whereas, these molecules have been shown to play a role in neurodegenerative diseases^(61)^, obesity, inflammation, diabetes and cancer^(62)^. These findings show that not all of the mechanisms regulating autophagy in other diseases are involved in the mutant COMP induced autophagy block in growth plate chondrocytes. Specific to this model system mechanism, increased mTORC1 signaling and suppression of SIRT1 reduces autophagosome maturation causing the autophagy blockade. Treatment of MT-COMP mice with resveratrol reactivates autophagy and cellular proliferation, while repressing intracellular retention and inflammation (**Fig. 3**).

The pleotropic benefits from resveratrol, shown in the mechanistic model depicted in **Fig. 4,** makes it uniquely effective in addressing the multifaceted mutant COMP pathology. Resveratrol diminishes cellular stresses including inflammation, oxidative stress and blocked autophagy resulting in anti-aging, anti-cancer, neuro-protective, anti-diabetic and cardio-protective properties)^(63–68)^. Resveratrol promoted autophagy in the mutant COMP pathology by upregulating SIRT1 which increased the number of LC3 positive autophagosomes (**Fig. 1** and **4B**). Resveratrol also reduced TNFα which results in repressing mTORC1 signaling through decreased MID1, pAKT, pS6 and increased PP2A^(4)^ (**Fig. 4**). Furthermore, treatment with resveratrol promotes autophagy by reducing oxidative stress and inflammation, which in turn alleviates ER stress^(4)^ (**Fig. 4**). These results demonstrate that resveratrol acts upon the mutant COMP pathology at multiple therapeutic points including ER stress, oxidative stress, inflammation (IL-1β; IL-16; OSM; FCF-L/YM1; TNFα/TRAIL), autophagy, chondrocyte death and proliferation ultimately resulting in a 50% recovery of loss of long bone growth (**Fig. 4**)^(3,4)^.

Although proteasomal degradation is not known to be the main mechanism clearing misfolded or aggregated protein from the ER, its role in a storage disease cannot be ruled out ^(48)^. Pharmacological blockade of proteasomal degradation along with resveratrol treatment resulted only in a minor decrease in clearance of mutant COMP (**Fig. 1**), indicating that proteasomal degradation is not the primary cellular mechanism by which resveratrol stimulates removal of mutant COMP from the ER of growth plate chondrocytes. Moreover, we demonstrated that retention of COMP in the ER allows for the assembly of an ordered matrix within ER vesicles of MT-COMP mice^(7)^. Once an intracellular matrix forms, there are no degradative enzymes to break up this large complex, which is likely too large to be translocated out of the ER to the proteasome for degradation^(13)^.

Importantly, the exciting findings from these studies show that there is a post-diagnosis window in which resveratrol diminishes growth plate pathology and clears mutant COMP from the ER in MT-COMP mice. Using our MT-COMP mouse that models PSACH clinical and molecular pathology in growth plate chondrocytes^(7)^, we showed that treatment with resveratrol from birth to adulthood successfully reduces murine growth plate pathology and restores some limb growth^(3)^, indicating that resveratrol may be a therapeutic for PSACH. Early resveratrol treatment could be implemented in familial cases where diagnosis of PSACH can be made at birth. However, 80% of PSACH cases occur *de novo*, therefore, diagnosis occurs later, between 18-24 months, when linear growth failure is recognized^(69,70)^. Here, we asked whether resveratrol treatment postdiagnosis would be effective for *de novo* cases. Resveratrol treatment beginning from birth or 1 week of age had the best outcomes, substantially reducing COMP retention, inflammation and cell death and initiating the recovery of DNA proliferation in growth plate chondrocytes. Approximating from human to mouse growth (1 week of mouse growth is roughly equivalent to 2 years in human growth^(71)^), our findings suggest that resveratrol treatment at or shortly after diagnosis will likely provide some reduction of clinical and growth plate pathology. Importantly, 3 weeks of resveratrol treatment (treatment from 1 - 4 wks) in the MT-COMP mouse increased autophagy (reduced pS6/mTORC1 signaling) demonstrating that autophagy reactivation is linked to the recovery of chondrocyte proliferation which most likely accounts for the recovery of some limb growth with resveratrol treatment. Interestingly, treatment beginning as late as 3 weeks of age (treatment from 3 - 4 wks) reduced inflammation and, therefore, would be expected to decrease joint pain suggesting even late treatment may have some benefit. The results for aspirin were similar to those with resveratrol with the exception that at least 2 weeks of aspirin treatment was required to reduce intracellular MT-COMP (**Supplementary Figs. 1** and **3**). Although aspirin reduced much of the mutant COMP growth plate pathology, aspirin is not approved for use in children under age 3 and caution must be used when administering aspirin to children/teens due to its association with Reyes syndrome (https://www.mayoclinic.org/diseases-conditions/reyes-syndrome/symptoms-causes/syc-20377255). Importantly, resveratrol was more effective than aspirin at reducing mutant COMP ER accumulation most likely related to its effect on multiple molecules that stimulate autophagy and reduce inflammation and oxidative stress (**Fig. 4**). A combination of aspirin and resveratrol treatment was not superior to resveratrol therapy alone in MT-COMP mice suggesting that aspirin and resveratrol mechanisms of action overlap (data not shown).

The outcome of these studies have important therapeutic applications because they show there is a therapeutic window during juvenile mouse growth in which resveratrol treatment diminished the mutant COMP chondrocyte pathology. This is a significant finding since most PSACH cases are not evident at birth. The absence of linear growth prompts diagnostic testing in children leading to PSACH diagnosis by 2 years of age. Moreover, these findings have important implications for addressing PSACH childhood pain that has not previously been appreciated or systematically treated^(5,16,72,73)^. Resveratrol’s therapeutic use in PSACH children will be tested in future clinical trials. Currently, a phase 1 clinical trial is underway to assess the efficacy of resveratrol and its potential to relieve adult PSACH joint pain (NCT03866200 clinicaltrials.gov), although, earlier treatment would likely be more effective based on the findings presented here.

Resveratrol treatment is promising given that the potential therapeutic window spans the post-diagnosis period in PSACH and resveratrol decreases multiple cellular stresses, stimulates autophagy clearance of MT-COMP allowing chondrocytes to function and proliferate potentially increasing long bone growth. Collectively, these findings suggest that chondrocyte death, inflammation and pain are likely to improve prior to recovery of growth with resveratrol therapy and earlier treatment results in the best outcome.

## Acknowledgments

We thank Frankie Chiu for technical assistance, dedication and organization. The authors have no conflict of interest to declare.

---

**Supplemental Figure 1:**
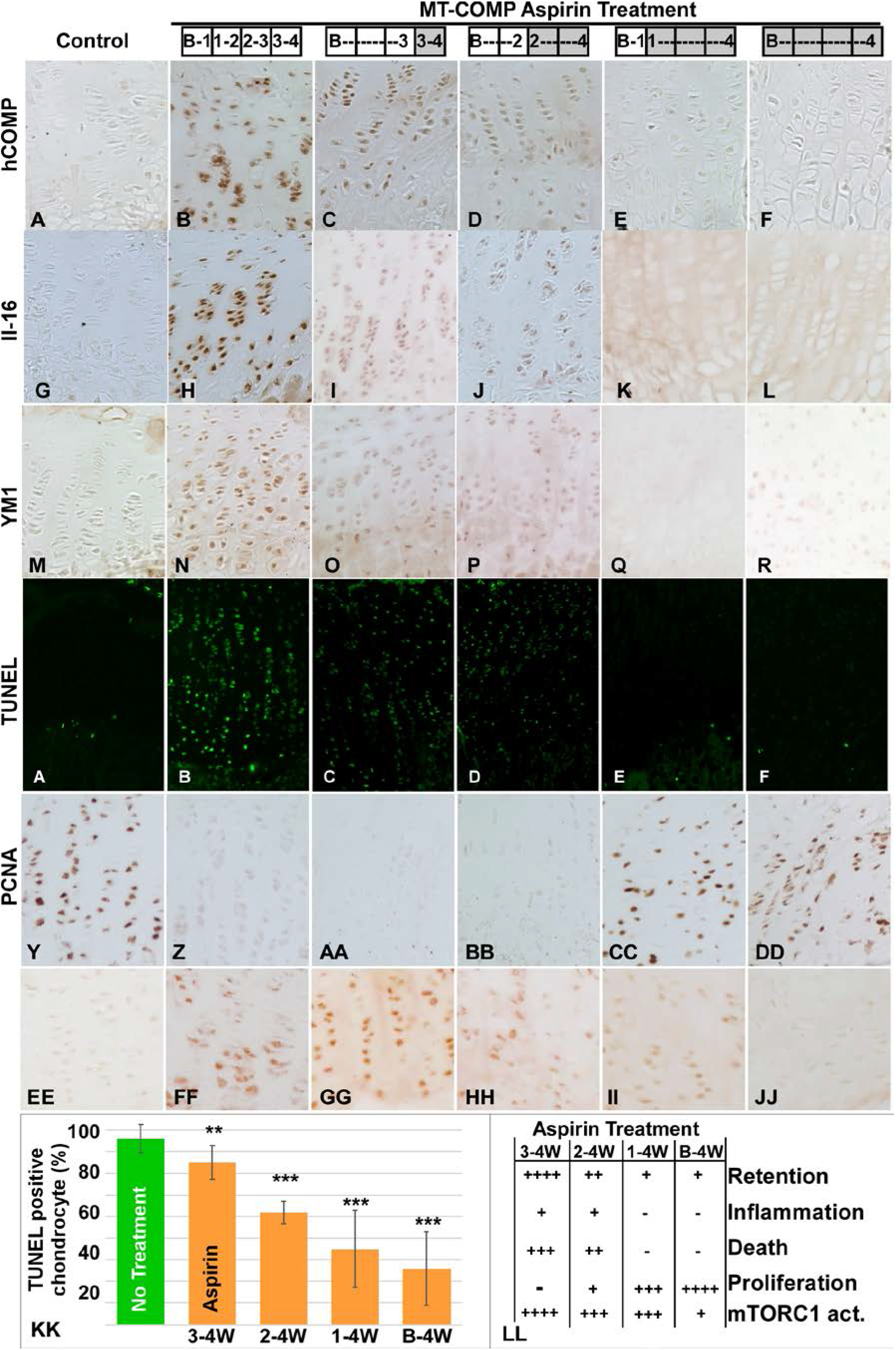
Aspirin treatment window. Treatments started at 1 or 2 or 3 weeks after birth and stopped at 4 weeks and evaluated for mutant COMP pathology. Each treatment group included 10 mice. Aspirin treatments are shown in the shaded boxes. Treatments starting at birth or 1 week of age produced the best therapeutic outcomes (**K, L**), later therapy mitigated but did not eliminate the disease progression (**C**). Control (C57BL\6) (**A, G, M, S, Y, EE**) and MT-COMP with no treatment (**B, H, N, T, Z, FF**) resveratrol treatment for 1 (**C, I, O, U, AA, GG**), 2 (**D, J, P, V, BB, HH**), 3 (**E, K, Q, W, CC, II**) or 4 (**F, L, R, X, DD, JJ**) weeks and growth plates at 4 weeks were evaluated for human COMP retention, IL-16 and YM1, TUNEL (cell death), PCNA and mTORC1 signaling. Quantification of TUNEL positive chondrocytes for different treatment periods are shown in **KK**. Bars = means with standard deviation. Aspirin treatment decreased intracellular MT-COMP after two weeks of treatment (**D**) and decreased in IL-16 and YM1 inflammation after one week of treatment (**I, O**) compared to (**H, N).** Aspirin decreased cell death after 2 weeks of treatment (**U**), mTORC1 signaling after 3 weeks of treatment (**JJ**) and increased proliferation after 3 weeks of treatment (**CC**). The relative change of MT-COMP pathology on treatment period is shown in **LL**.

**Supplemental Figure 2.**
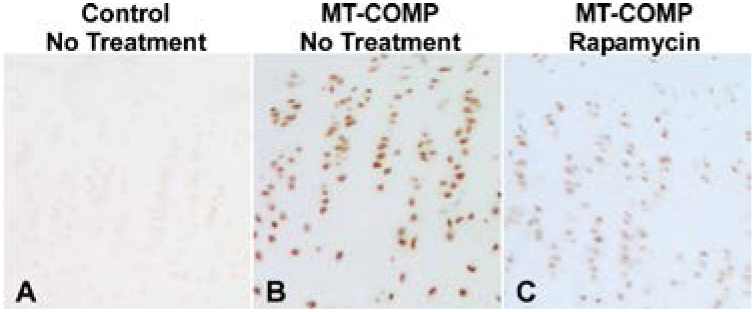
Rapamycin activates autophagy in MT-COMP growth plate chondrocytes. Rapamycin reduces MT-COMP retention in chondrocytes. Immunostaining with human COMP antibody of: **A** MT-COMP growth plate chondrocytes at 4 weeks with no treatment**, B** control vehicle, **C** rapamycin (1 mg/kg 5 IP injections per week from 1 to 4 weeks of age). Ten mice were included in each treatment group.

